# Microtubules provide a viscoelastic resistance to myocyte motion

**DOI:** 10.1101/361824

**Authors:** M. A. Caporizzo, C. Y. Chen, A. K. Salomon, K. Bedi, K. B. Margulies, B. L. Prosser

## Abstract

**Background:** Microtubules (MT) buckle and bear load during myocyte contraction, a behavior enhanced by post-translational detyrosination. This buckling suggests a spring-like resistance against myocyte shortening, which could store energy and aid myocyte relaxation. Despite this visual suggesting of elastic behavior, the precise mechanical contribution of the cardiac MT network remains to be defined.

**Methods:** Here we experimentally and computationally probe the mechanical contribution of stable microtubules and their influence on myocyte function. We use multiple approaches to interrogate viscoelasticity and cell shortening in primary murine myocytes where either MTs are depolymerized or detyrosination is suppressed, and use the results to inform a mathematical model of myocyte viscoelasticity.

**Results:** MT ablation by colchicine concurrently enhances both the degree of shortening and speed of relaxation, a finding inconsistent with simple spring-like microtubule behavior, and suggestive of a viscoelastic mechanism. Axial stretch and transverse indentation confirm that microtubules increase myocyte viscoelasticity. Specifically, increasing the rate of strain amplifies the MT contribution to myocyte stiffness. Suppressing MT detyrosination with parthenolide or via overexpression of tubulin tyrosine ligase (TTL) has mechanical consequences that closely resemble colchicine, suggesting that the mechanical impact of MTs relies on a detyrosination-dependent linkage with the myocyte cytoskeleton. Mathematical modeling affirms that alterations in cell shortening conferred by either MT destabilization or tyrosination can be attributed to internal changes in myocyte viscoelasticity.

**Conclusions:** The results suggest that the cardiac MT network regulates contractile amplitudes and kinetics by acting as a cytoskeletal shock-absorber, whereby MTs provide breakable cross-links between the sarcomeric and non-sarcomeric cytoskeleton that resist rapid length changes during both shortening and stretch.

## 1. Introduction

The proliferation of microtubules (MT) and their post-translational detyrosination correlates with declining cardiac function{Fassett:20l3gi, Robison:2016bc, Sato:1997tw, Chen:20l8dx}. Recently, we observed that MTs buckle in a sinusoidal fashion during myocyte contraction, suggesting a spring-like mechanism that could resist compression and possibly provide restoring force to facilitate cardiac relaxation. MT buckling relies on post-translational detyrosination (dTyr) (for review of the tyrosination cycle, see(4)), which facilitates interaction between MTs and the cardiac sarcomere(2) and is associated with MT stability(5).

While MT buckling provides visual evidence suggestive of an elastic resistance to contraction, previous efforts to pin down the mechanical contribution of MTs have produced inconsistent results. Magnetic twisting cytometry and single cell shear measurements show MTs provide both viscous and elastic rigidity to the cardiomyocyte(6-8). In contrast, measurements of steady state elasticity by myocyte stretch have shown a small(9) or insignificant(6) role of the MT network in axial stiffness. Multiple groups have observed increased contractility with MT destabilization(10-12), which is most often attributed to decreased mechanical resistance, but this finding has not been supported by all(13).

Altered calcium handling must also be considered, as MTs mediate the mechanotransduction of calcium signals (14, 15) and the transport of proteins required for normal excitation-contraction (EC) coupling (16, 17). Pharmaceutical ablation of MTs was reported to increase the calcium transient during excitation-contraction coupling(18), but this has been challenged by multiple groups who showed no effect of MT destabilization on EC coupling (15, 19). Recent work showed that reducing MT-dTyr similarly enhances myocyte contractility and reduces myocytes stiffness without grossly affecting EC coupling (2). In diseased myocytes, where a proliferated MT network likely plays a more significant mechanical role, MT destabilization has been shown to enhance myocyte contractility independent of a change in the calcium transient {Zile:1999wu, Chen:2018dx}. Thus, despite apparent conflicting results, increased mechanical compliance remains a sensible explanation for the enhanced contractility observed upon MT destabilization, but it remains to be clearly demonstrated how an enhancement in contractility may arise from a reduction in MT-dependent stiffness.

This study aims to dissect the precise role that MTs and MT-dTyr play in cardiomyocyte viscoelasticity, and to determine if altered viscoelasticity is sufficient to explain changes in myocyte contraction. To this end we paired calcium and contractility measurements with high-sensitivity viscoelastic analysis using transverse nanoindentation and tensile stress-relaxation tests. We find that MT tyrosination or depolymerization similarly amplifies and accelerates contractility independent of a change in calcium handling. Viscoelastic measurements that inform computational modeling suggest these changes can be completely accounted for by measured reductions in myocyte viscoelasticity while they cannot be explained by changes in passive stiffness or elasticity alone.

## 2. Methods

This study was conducted on freshly isolated adult ventricular myocytes from Sprague-Dawley rats. All experiments were performed immediately following cell isolation with the exception of adenoviral treated myocytes.

### 2.1 Murine models

All animal care and experimental procedures were approved and performed in accordance with the ethical standards set forth by the Institutional Animal Care and Use Committee of the University of Pennsylvania and by the National Institutes of Health. Rats (Sprague Dawley) were purchased from Harlan labs.

### 2.2 Myocyte Isolation and culture

Primary adult ventricular myocytes were isolated from 8-12 week old Sprague-Dawley rats as previously described (14). In summary, rats anesthetized under isoflurane received an intramuscular injection of heparin (~1000 units/kg) before the heart was excised and cannulated for retrograde perfusion with enzymatic digestion solution at 37^o^C on an Langendorff apparatus. Digested hearts were then minced and triturated with glass pipettes to isolate individual cardiomyocytes. Isolated myocytes were gently centrifuged (300 RPM) to remove debris and gradually reintroduced into rat cardiomyocyte (RCM) medium (formulation below).

Following isolation, cells in RCM medium with 25 μM cytochalasin D were plated at low density into 12 well plates and treated with viral constructs. Viral constructs were expressed for 48 h with MOI = 100-200 of adenoviral TTL-IRES-dsRed as previously described(2), and experiments were conducted on the second day after isolation. Freshly isolated myocytes were diluted 5x from RCM medium into normal Tyrode’s solution (NT) and treated with 10 μM of either colchicine or parthenolide. Experiments were conducted after 1.5 h drug treatment in the presence of 10 μM colchicine or parthenolide. All experiments and culture on rat cardiomyocytes were conducted in the presence of 1.8 mM Ca^2+^.

RCM Medium: Medium 199 (GIBCO, 11150-59) was supplemented with 1x Insulin-Transferrin-Selenium-X (GIBCO 51500-56), 20 mM HEPES at pH 7.4, and 0.1 mg/ml Primocin.

Normal Tyrode’s: 140 mM NaCl, 0,5 mM MgCl2, 0.33 mM NaH2PO4, 5 mM HEPES at pH 7.4, 5.5 mM glucose, 5 mM KCl, titrated to a pH of 7.4 with 1 M NaOH.

### 2.3 Cell Stretch

Experiments were performed in custom-fabricated glass bottom petri dishes mounted on an LSM Zeiss 880 inverted confocal microscope using a 40x oil 1.4 NA objective and transmitted light camera (IonOptix MyoCam-S). Cell stretch experiments were carried out as previously described(2). Cells were attached to glass holders with a laser-etched cavity via MyoTak (Ionoptix) adhesive. One cell holder was connected to a piezoelectric length controller and the other to a high-sensitivity optical force transducer (IonOptix OFT-100). Force and sarcomere length were continuously recorded at 1 kHz. Cells with a resting sarcomere length below 1.7 μm were considered hypercontracted and discarded from these studies. Force and sarcomere length traces were analyzed in IonWizard (IonOptix).

### 2.4 Calcium and Contractility Measurements

Myocytes were loaded by 15 min incubation at 1 uM Fluo-4-acetoxymethyl ester (Invitrogen). Imaging was conducted in confocal line scan mode with a 488-nm argon ion laser at 0.909 ms/line. Cells were electrically paced (IonOptix MYP100, IonWizard) at 1 Hz for 20 s to achieve steady state; the final 5 traces of the pacing protocol were pooled and analyzed for calcium transient properties. The measured fluorescence (F) throughout the transient was normalized to the resting fluorescence prior to stimulation (F_0_) to normalize for heterogeneity in dye loading. Simultaneously, sarcomere length was measured optically by Fourier transform analysis (IonWizard, IonOptix) and during 20 s of pacing, the final five traces were again recorded and averaged using IonWizard.

### 2.5 Nanoindentation

Mechanical properties at the microscopic scale were measured using nanoindentation (Piuma, Optics11, The Netherlands). Freshly isolated human myocytes were attached to glass bottom dishes coated with MyoTak (IonOptix) in NT solution at 1 mM Ca^2+^. A spherical indentation probe with a radius of 3.05 μm and a stiffness of 0.026 N/m was used. Myocytes were indented to a depth of 2.5-3.5 μm with velocities of 0.1, 0.25, 0.5, 1.0, 2.0, 5.0, 10.0, 20.0, 50.0, 100.0, and 150.0 μm/s. The tip was held in this indentation depth for 1 s, and retracted over 2 s. The Young’s moduli were calculated automatically by the software by fitting the force vs. indentation curve to the Hertz equation. The Young’s modulus E is derived from the fit of the initial 60% of the loading force-displacement curve (F(h)), the indenter tip radius (R) and indentation depth (h), according to the following formula, for which a Poisson’s ratio (ν) of 0.5 was assumed.

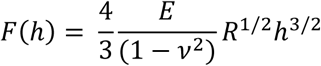

Average of E in each condition with standard error is plotted against different indentation speed. Low velocity indentation measures elastic contributions (E_min_) to stiffness, high velocity indentation measures elastic and viscous contributions (E_max_). The change in modulus with rate represents myocyte viscoelasticity (ΔE). To fit the elastic modulus vs. indentation-rate plots to

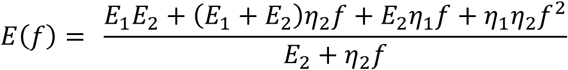

the VE model, indentation velocity was converted to indentation frequency using the average indentation depth at each rate. The viscoelastic model could then be solved as a function in indentation frequency allowing the viscoelastic constants E_1_, E_2_, η_1_ and η_2_ to be determined from the fit of the expression below.

### 2.7 Statistical Analysis

Statistical analysis and graphing were performed with OriginLab 9.0. Values are presented as means ± standard error in bar and line graphs. Dot plots show standard deviation as whiskers and mean line. Where comparisons between sets were both repetitive and restricted, the Bonferroni multiple comparisons correction was used to adjust the significance threshold of two-sided T-tests accordingly. One-way ANOVA with post-hoc Tukey test was used to correct for multiple comparisons sharing a single control condition.

### 2.8 Mathematical Model for myocyte contractility

For stress exerted by the contractile unit during the contractile timescale, σ(t), the strain response, ε(t)=ΔSL/SL_o_(t), can be analytically determined by solving the strain response of the VE model proposed in Figure 2D, which is a Kelvin-Voigt viscoelastic element in parallel configuration with a Maxwell element. The total stress exerted by the sarcomeric units is distributed equally over the parallel units in the model by their corresponding stress-strain relationship.

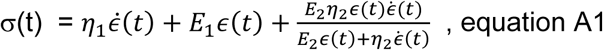

Simplifying the expression for stress and strain yields the second order ordinary differential equation below.

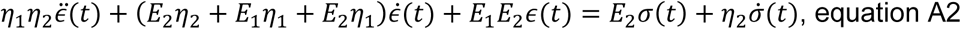

This is a second order ODE of the form:

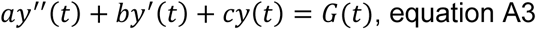

Where:

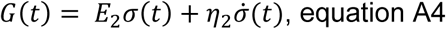

Thus, for an arbitrary input stress, G(t) can be determined. The solution for *ϵ*(*t*) to the ordinary differential equation (equation A3) is written explicitly in terms of elastic elements as follows:

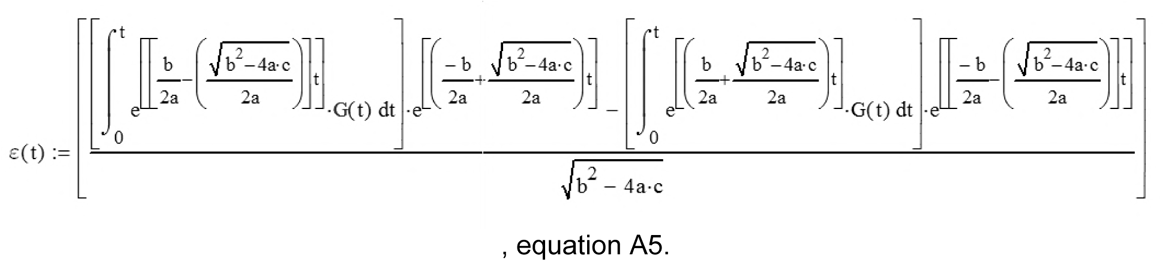

where,

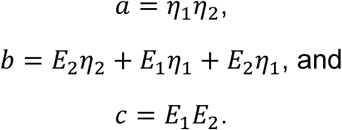

The limitations of the model require that the solution be real, i.e. b^2^ > 4ac, and solvable, i.e. a > 0. Consequently, the solution is valid only when each element has a finite positive value. This is reasonable considering we are interested in modelling the case where all of the elements have a contribution to the mechanics. Further, in order to estimate the timescale of the force relative to the calcium transient, we chose the function (Figure 5 A) which has a shape (rise and decay parameters) that we felt was consistent with the force that would be generated by the measured calcium transients.

Wolfram Alpha was used to determine the solution for the ODE, and Mathcad 14.0 (PTC Inc.) was used to generate the analytical model relating the viscoelastic parameters to contractility (strain, ε(t)).

## 3. Results

### 3.1 Depolymerizing MTs enhances contractile and relaxation velocity

We collected paired calcium and contractility transients on rat cardiomyocytes (RCMs) treated with colchicine (Figure 1) immediately following cell isolation. As shown in Figure 1A, the calcium transient is largely unaffected by treatment with colchicine (purple). Peak calcium, rise times, and decay times are not significantly different in colchicine treated cells (Figure 1B). In contrast, myocyte contractility is enhanced by colchicine (Figure 1C). Resting sarcomere length is unchanged (~1.83um, Table S1), but fractional shortening is significantly increased. The increased fractional shortening occurs over a similar timescale (Figure 1D), leading to higher peak contractile and relaxation velocities for colchicine treated cells (Figure 1F). Thus, colchicine can increase the contractility of RCMs independent of an increase in [Ca^2+^]_i_, suggesting that MT depolymerization affects the rate at which myocytes can change their length under a similar level of activating [Ca^2+^]_i_.

**Figure 1:**
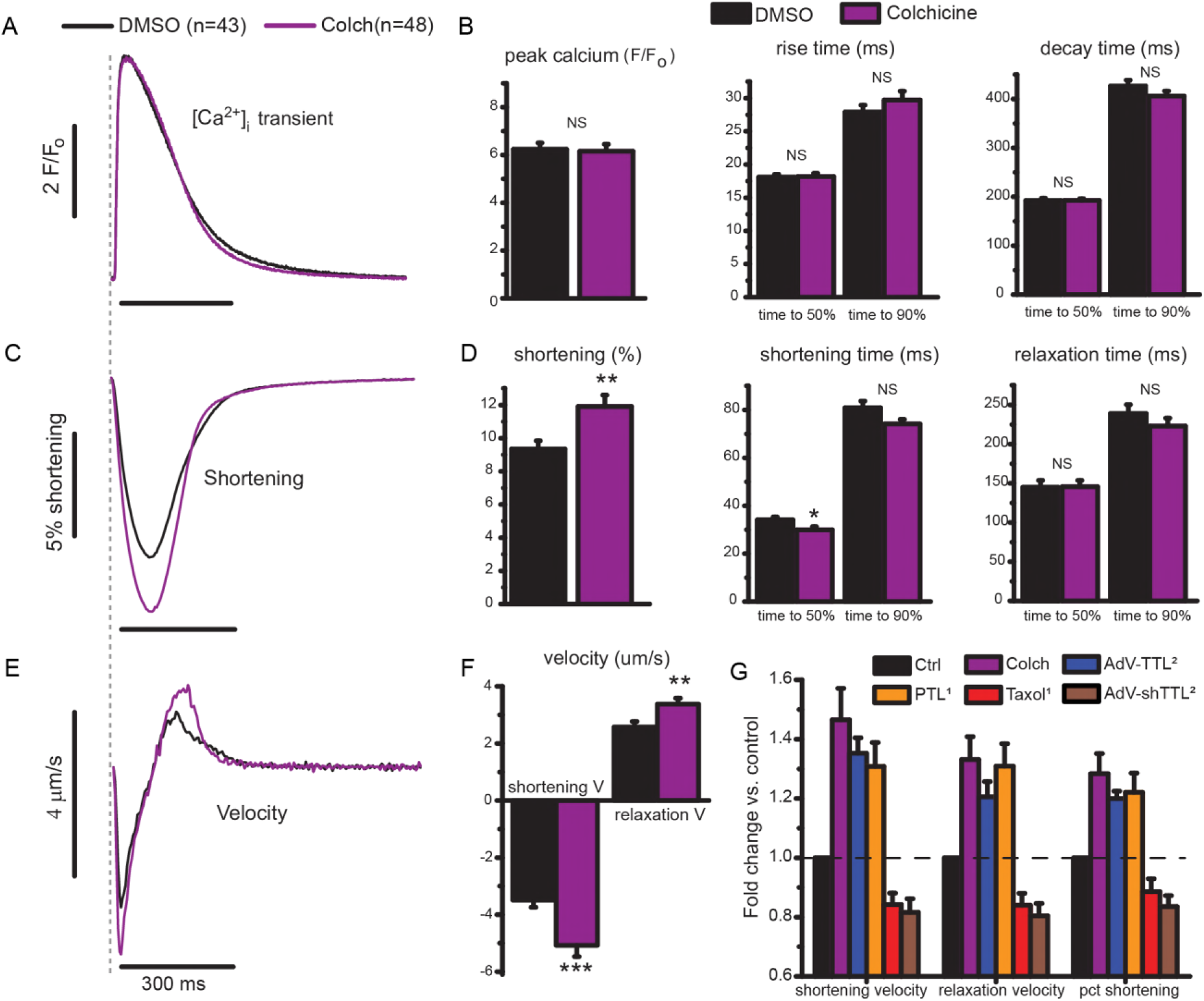
MT depolymerization enhances myocyte contractility. **A,** Normalized calcium transient (F/F_o_) for control (black) and colchicine (purple) treated cells. **B**, Peak calcium, time to peak, and decay time determined from individual traces (N=6 hearts, n=43 DMSO cells, n=48 colchicine treated cells.) **C**, Relative shortening (change in sarcomere length divided by resting sarcomere length) with time. **D**, Peak shortening, shortening and relaxation times from individual traces from identical cells analyzed in B. **E**, Average shortening velocity for DMSO and colchicine treated RCMs from **A** and **C**. **F**, Peak shortening and relaxation velocity determined for individual data points. Statistical significance was determined as ^⋆⋆^p<0.01, or ^⋆⋆^p<0.001 compared to DMSO, student t-test. **G,** Comparison of changes in contractile velocities and fractional shortening upon various manipulations to depolymerize MTs or alter MT detyrosination. Each treatment condition is displayed normalized to its relative control (for example DMSO treated for PTL, or a null-encoding adenovirus (AdV-Null) for AdV-TTL). Colchicine data is from the current study. ^1^ PTL and taxol data are replotted from(l5). ^2^TTL overexpression and knock down data are re-plotted from (2). Data in B, D, F and G are means with standard error bars.

A quantitatively similar calcium-independent enhancement of contractility has been observed when post-translational detyrosination of the MT network is suppressed (Figure 1G)(2, 15). This system can be pushed in both directions, as myocyte contraction is reduced and prolonged when MT density or detyrosination is increased (Figure 1G). If mechanical in origin, the concurrent increase in contractile and relaxation velocity associated with a reduction in MTs or MT-dTyr points towards a viscous or viscoelastic role for dTyr MTs. This is in contrast to a purely elastic mechanical role where an increased fractional shortening would accompany a prolonged contractile timescale. Thus, our data here and in recent work predicts a viscous role for dTyr MTs in resisting myocyte length change.

### 3.2 Depolymerizing or tyrosinating MTs similarly reduces myocyte viscoelasticity

To determine the contribution of stable MTs to cardiomyocyte mechanical properties, transverse indentation was performed at variable speeds to obtain a viscoelastic profile of the cell and to extract generalized viscoelastic parameters (21). Figure 2A and B show the elastic modulus as a function of indentation velocity. The increase in myocyte stiffness with indentation velocity confirms that cardiomyocytes are viscoelastic. Further, the data can be used to derive a generalized viscoelastic (VE) model (fit by solid lines) for which the simplest model that fits the data is depicted in Figure 2D, consisting of a spring, dashpot, and Maxwell element in parallel. The spring E_1_ describes the steady state stiffness, the spring E_2_ becomes engaged at timescales faster than η_2_/E_2_ but dissipates stored energy if compressed for longer, and the viscous element η_1_ approximates cytoplasmic viscosity which accounts for the upward trend of modulus at rates > 50 μm/s.

**Figure 2:**
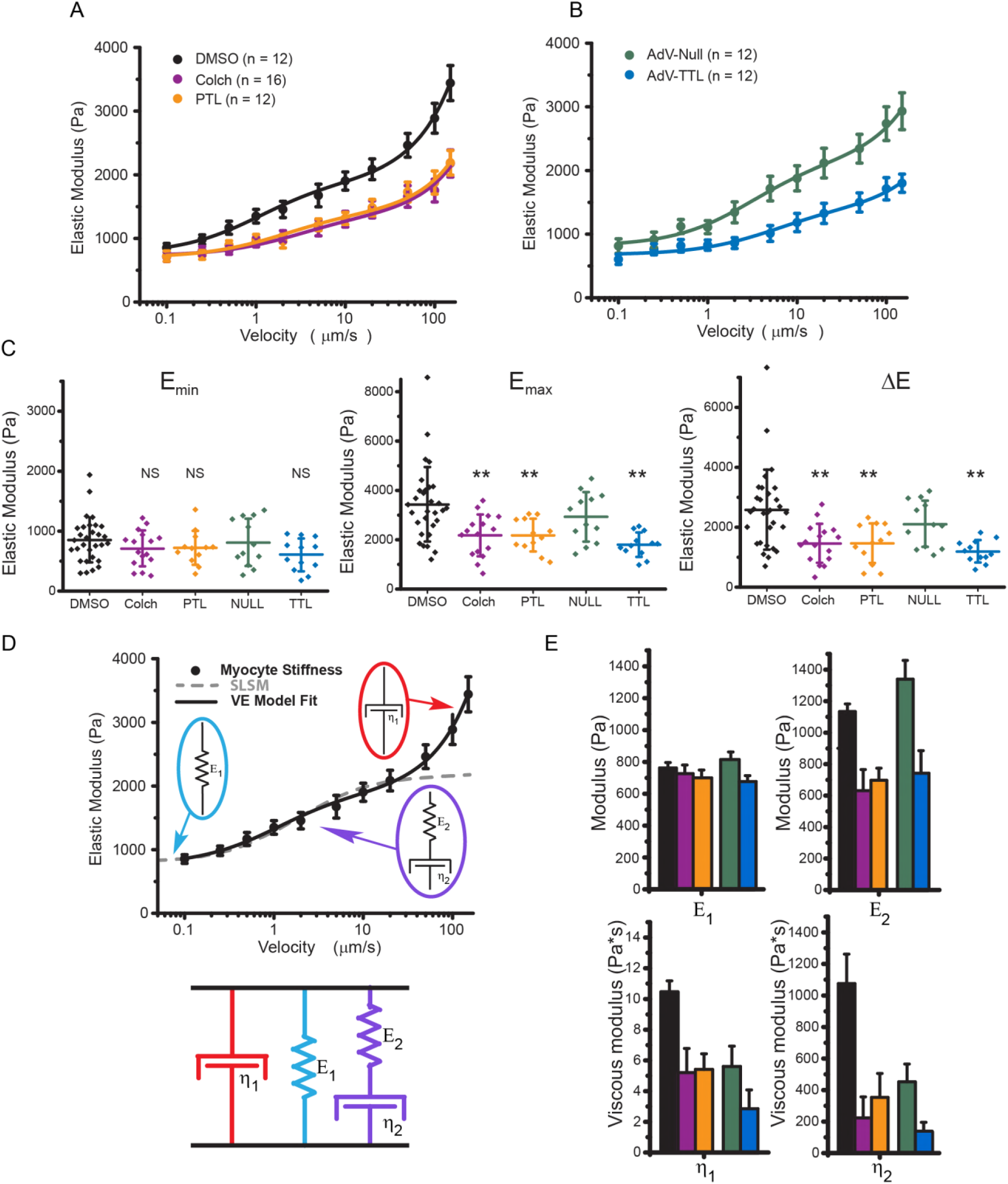
Transverse indentation at variable rates to assess myocyte viscoelasticity. **A**, Elastic modulus vs. indentation velocity in freshly isolated RCMs treated with DMSO, colchicine, or PTL. Data points are means with standard error whiskers. (n=12, 16, and 12, cells from 3 rats for DMSO, colchicine, and PTL respectively.) **B**, Elastic modulus vs. indentation velocity in RCMs 48h post transfection with adenovirus Null, green, or AdV-TTL, blue. Data points are means with standard error whiskers. (n=12 cells from 3 rats for both Null and TTL) Solid lines in both A and B are best fit curves to VE model shown in **D**. **C**, The minimum elastic modulus, E_min_, determined by the cell stiffness at the slowest (100 nm/s) indentation rate. The maximum elastic modulus, E_max_, is determined by the cell stiffness at the fastest (150 p,m/s) indentation rate. The change in elastic modulus over the velocity range measured, ΔE, is determined by E_max_-E_min_ for each cell. Statistical significance was determined by one-way ANOVA with posthoc Tukey as ^⋆^p<0.05, ^⋆⋆^p<0.01, or ^⋆⋆⋆^p<0.001 compared to DMSO or Null. Dot plots show mean line and whiskers as standard deviation. **D,** The VE model (a Maxwell element in parallel configuration with a Voigt element) for which the velocity-dependence of the Elastic modulus can be fit to extract the stiffness of each element. **E,** Viscoelastic parameters extracted from model fit to data in **A** and **B.** Bar plots show the estimated fitted value with whiskers showing the fit certainty.

Colchicine treated cardiomyocytes are still viscoelastic and exhibit similar steady state stiffness (E_min_, Figure 2C) as untreated cells. The stiffness of colchicine treated cells increases less with indentation rate, indicating a reduction in viscoelasticity (ΔE, Figure 2C). The difference in stiffness becomes significant at indentation rates > 500 nm/s and the total change in modulus over the velocity range tested (ΔE) is significantly less for colchicine treated cells (Figure 2E). Using the VE model to derive the stiffness of individual elements, the steady state elasticity, E_1_, is not different for colchicine treated cells, while viscoelastic parameters, E_2_, η_1_, and η_2_ are all significantly reduced (Figure 2E). This is consistent with the MT network engaging with the surrounding structure in the myocyte through cross-links(2) that slip (and thus do not contribute to mechanical properties) on a timescale approximated by η_2_/E_2_, or ~ 1 s.

Previous work suggested a decrease in viscoelasticity upon MT tyrosination by TTL overexpression, but restricted indentation depth (<500 nm) limited measurement sensitivity to sub-membrane cortical stiffness (2). Thus, indentation of rat cardiomyocytes treated with PTL or overexpressing TTL are here performed at greater indentation depth (3 μm). A decrease in viscoelasticity nearly identical to that of colchicine treatment is observed when MTs are tyrosinated, whereby E_min_ is not different compared to the respective control while E_max_ and AE are significantly decreased (Figure 2C,E). No significant differences in stiffness between colchicine and PTL treated myocytes are observed, which is consistent with a detyrosination-dependent engagement of MTs with the sarcomeric cytoskeleton forming the basis of their contribution to myocyte viscoelasticity.

Recognizing that myocytes are anisotropic and nanoindentation probes only the transverse axis, single cell tensile tests (axial stretch) were conducted to determine the mechanical contribution of the MT cytoskeleton along the contractile axis. Immediately following isolation, we stretched myocytes at different rates to determine the contribution of strain rate to stiffness, and viscoelasticity was determined by a 5 second isometric hold to evaluate stress relaxation. Figure 3 shows the position of the length controller (top row), the corresponding change in sarcomere length applied to the cell (ΔSL, second row), and the measured force recorded (third row) as a function of time. In the 100 μm/s stretch the force transient exhibits a peak followed by a rapid decay (τ = 200 ms) and then a slow relaxation towards a steady-state value. The peak force and the magnitude of the relaxation (Figure 3 A, C) is reduced by both colchicine and PTL treatment, indicating a reduction in viscoelasticity. In contrast, in 2 μm/s stretches, significant stress relaxation is not observed irrespective of treatment group, and the relaxation during the isometric hold is not significant or affected by colchicine or PTL.

**Figure 3:**
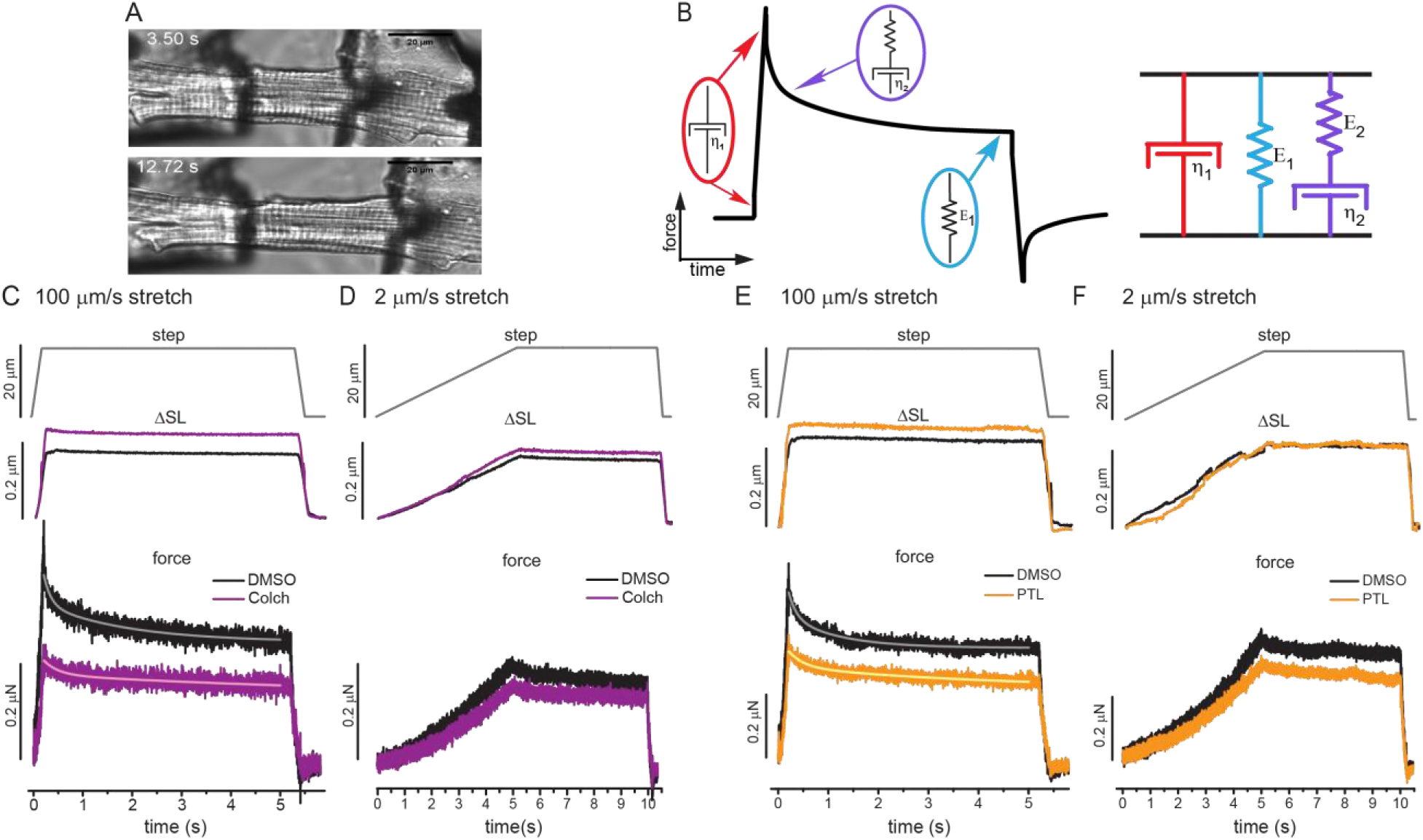
Axial stretch at variable rates following colchicine and parthenolide treatment of RCMs. **(A)** Trasmitted light image of isolated adult rat cardiomyocyte before and during tensile test. (B) The VE model (a Maxwell element in parallel configuration with a Voigt element) for which the time dependence of the stress-strain curve can be used to extract values for each parameter. Top row shows the imposed length change – a 20 μm length controller step over either a 200 ms **(C, E)** or 5 s **(D, F)** stretch, followed by a 5 s hold, and 200 ms return. Middle row shows the average change in sarcomere length in response to stretch. Bottom row shows the average change in force in response to stretch. **A-B,** 100 μm/s stretch and 2 μm/s stretch for DMSO and colchicine (N=5 hearts, n=17 cells DMSO, n= 18 cells colchicine). **C-D**, 100 μm/s stretch and 2 p,m/s stretch for DMSO, and PTL. (N=5 hearts, n=14 cells DMSO, n= 17 cells colchicine). Solid traces overlaid on force-relaxation curves are best fits to double exponential decay function.

Cardiomyocyte elastic modulus at each strain rate was determined from the slope of the stress (force divided by myocyte cross sectional area) versus strain (ΔSL/resting SL) curve (Figure 4). The area between the stretch (dark traces) and return (light traces) is an approximation of the amount of energy (work) lost to viscosity during each stretch. The average elastic modulus of each condition is indicated by the slope of the stress-strain curves, and peak and steady state modulus for individual myocytes is presented in Figure 4C. The peak modulus is reduced by colchicine and PTL for the fast (100 μm/s) but not the slow (2μm/s) stretch rate (Figure 4 A,B). The energy lost to viscosity is significantly reduced by colchicine and PTL treatment under rapid stretch, while the slow stretch (which shows minimal energy dissipation and almost purely elastic behavior) is not significantly altered by colchicine or PTL. Thus, both tensile tests and transverse indentation indicate that colchicine and PTL treatment reduce myocyte viscoelasticity, causing stiffness differences to be most pronounced at high strain rates. These findings are consistent with the contractility data in Figure 1, which suggests a viscoelastic change in the cardiomyocyte because of concurrent increases in contractile amplitude and velocity that cannot be explained by differences in the calcium transient.

**Figure 4:**
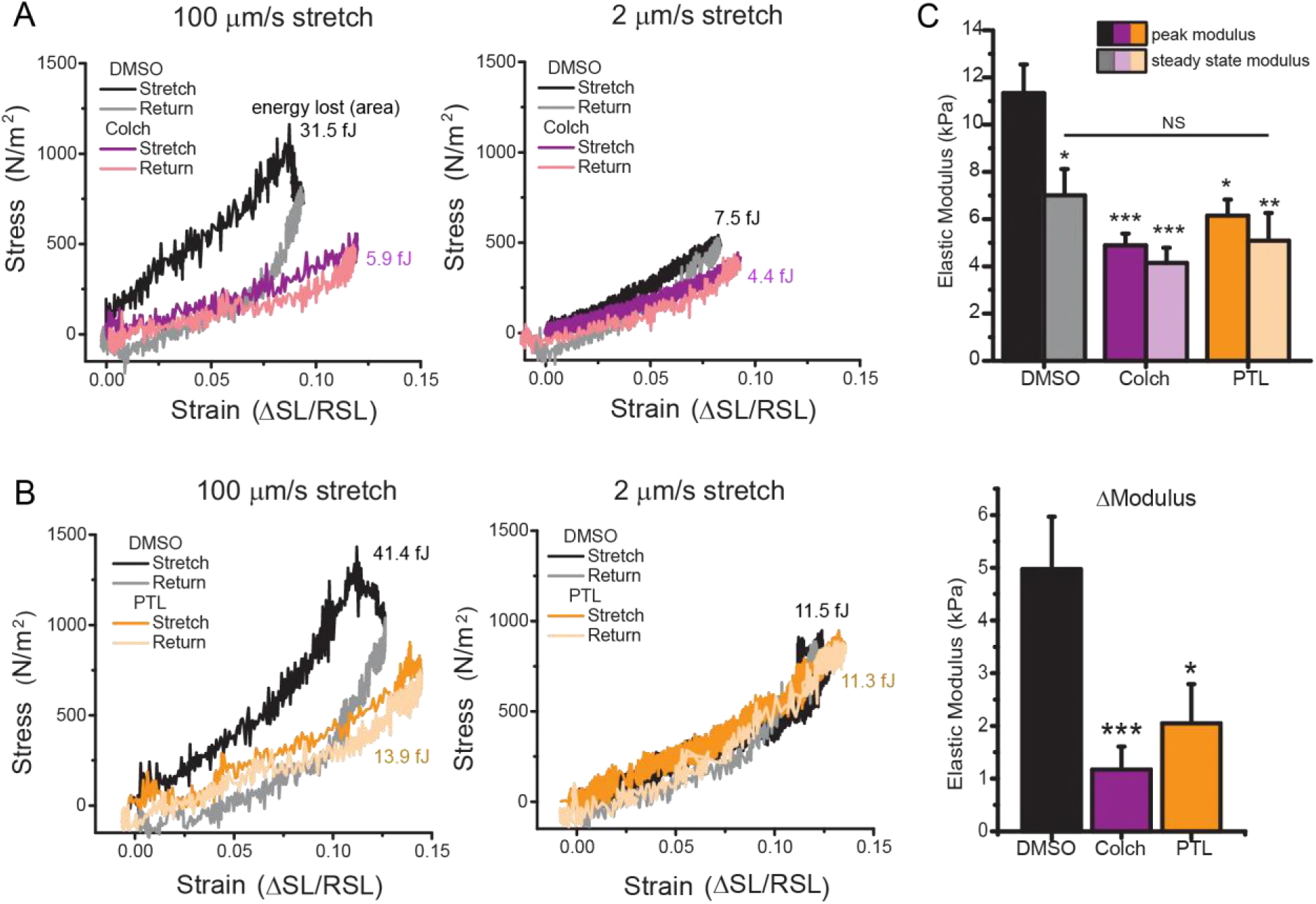
Stress-strain curves calculated from axial stretch. **A**, Stress-strain loops for DMSO and colchicine treated cells at 100 μm/s and 2 μm/s. **B,** Stress-strain loops for DMSO and PTL treated cells at 100 μm/s and 2 μm/s. **C**, Peak and steady state modulus (top), and relaxation modulus (peak – steady state) (bottom) for individual cells treated with DMSO, PTL, or colchicine. Statistical significance was determined by one-way ANOVA with post-hoc Tukey as ^⋆^p<0.05, ^⋆⋆^p<0.01, or ^⋆⋆⋆^p<0.001 compared to DMSO.

The VE model derived from transverse indentation data (Figure 2) can also be fit to the axial stretch data (Figures 3 and 4) to extract viscoelastic parameters along the contractile axis (methods). Table 1 shows a comparison of viscoelastic parameters derived from the two orthogonal approaches. Generally, there are stiffer elastic components, a higher viscosity and a faster relaxation time (τ=η_2_/E_2_) in the longitudinal vs. transverse axis of the cell. This is consistent with an elastic spring element, such as titin, and a viscoelastic element, such as the passive resistance of the myofilaments, contributing more to the mechanical properties of the myocyte during axial stretch but not upon transverse indentation.

**Table 1A:**
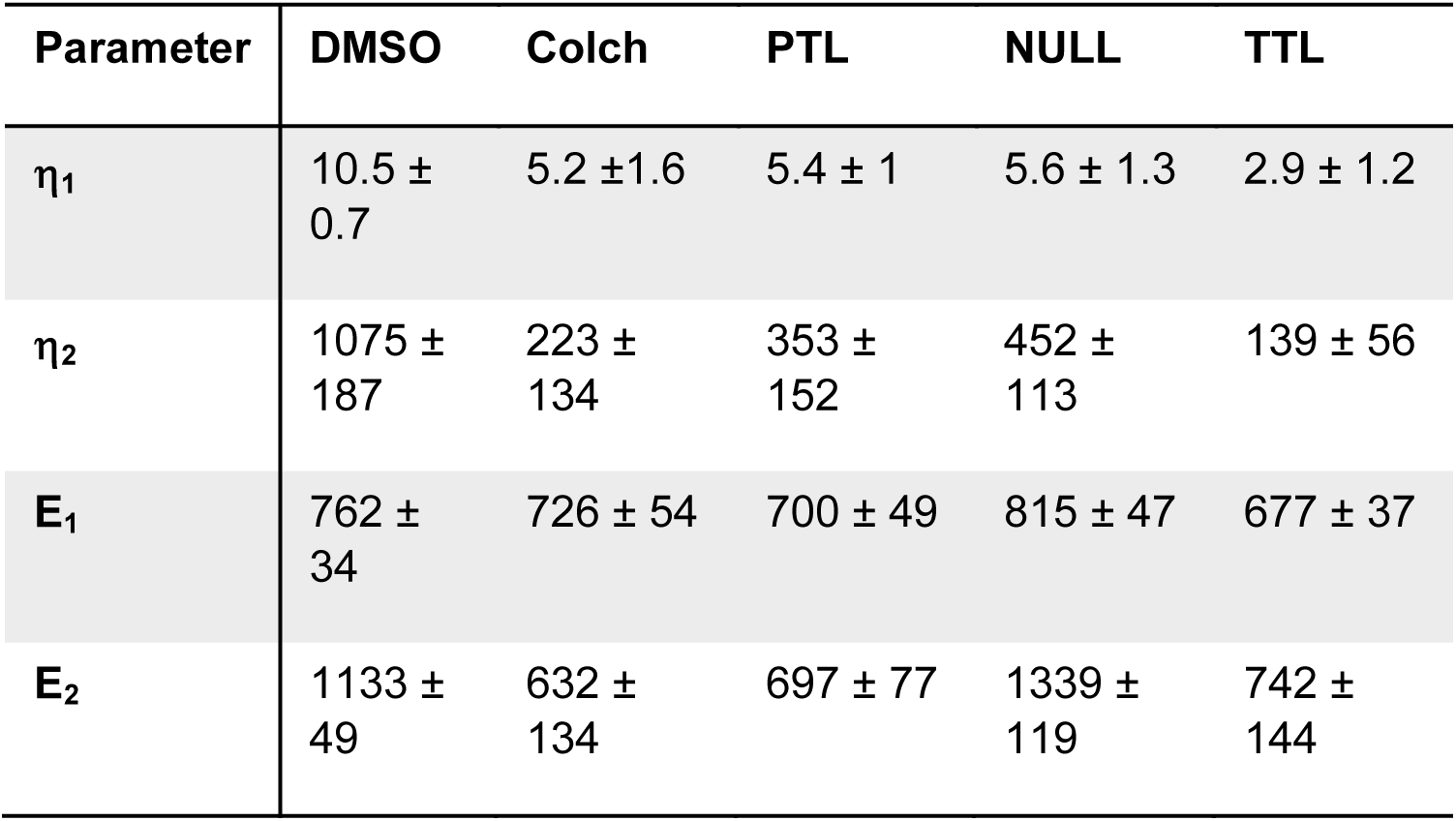
Viscoelastic parameters derived from transverse indentation. Values are expressed as mean ± fit error.

| Parameter | DMSO | Colch | PTL | NULL | TTL |
| --- | --- | --- | --- | --- | --- |
| $\eta_1$ | 10.5 ± 0.7 | 5.2 ± 1.6 | 5.4 ± 1 | 5.6 ± 1.3 | 2.9 ± 1.2 |
| $\eta_2$ | 1075 ± 187 | 223 ± 134 | 353 ± 152 | 452 ± 113 | 139 ± 56 |
| $E_1$ | 762 ± 34 | 726 ± 54 | 700 ± 49 | 815 ± 47 | 677 ± 37 |
| $E_2$ | 1133 ± 49 | 632 ± 134 | 697 ± 77 | 1339 ± 119 | 742 ± 144 |

**Table 1B:** Viscoelastic parameters derived from passive length tension. Elasticities are expressed as mean ± standard error, and viscosities are expressed as mean ± fit error.

| Parameter | DMSO | Colch | PTL | NULL | TTL |
| --- | --- | --- | --- | --- | --- |
| $\eta_1$ | 164 ± 15 | 73 ± 4 | 91 ± 12 | xx | xx |
| $\eta_2$ | 827 ± 193 | 434 ± 155 | 736 ± 232 | xx | xx |
| $E_1$ | 7108 ± 1111 | 4139 ± 644 | 5087 ± 1177 | xx | xx |
| $E_2$ | 4972 ± 998 | 1172 ± 434 | 2051 ± 743 | xx | xx |

### 3.3 A viscoelastic model for contractility

We next generated a computational model to test if the observed changes in viscoelasticity can account for enhanced myocyte contractility upon MT destabilization (Figure 1). Myocyte shortening (strain) was derived from the VE model (Figure 2D) to generate an analytical model that relates myofilament force production (internal stress) to myocyte length change (Figure 5). Force production was approximated by the function shown in Figure 5A for which the amplitude and decay constant, τ, determine the size and decay rate of the force. The force function is consistent with the shape and time-course of developed tension in cardiac muscle (22). The identical force (amplitude and τ) was used throughout Figure 5, based on the observation of similar Ca^2+^ transients (Figure 1) and assumption of similar myofilament Ca^2+^ sensitivity. Adjusting the stiffness of the viscoelastic elements in the model to fit (dashed line) the contractility trace (black solid line) indicates that the VE model can accurately capture the contractile transient.

**Figure 5.**
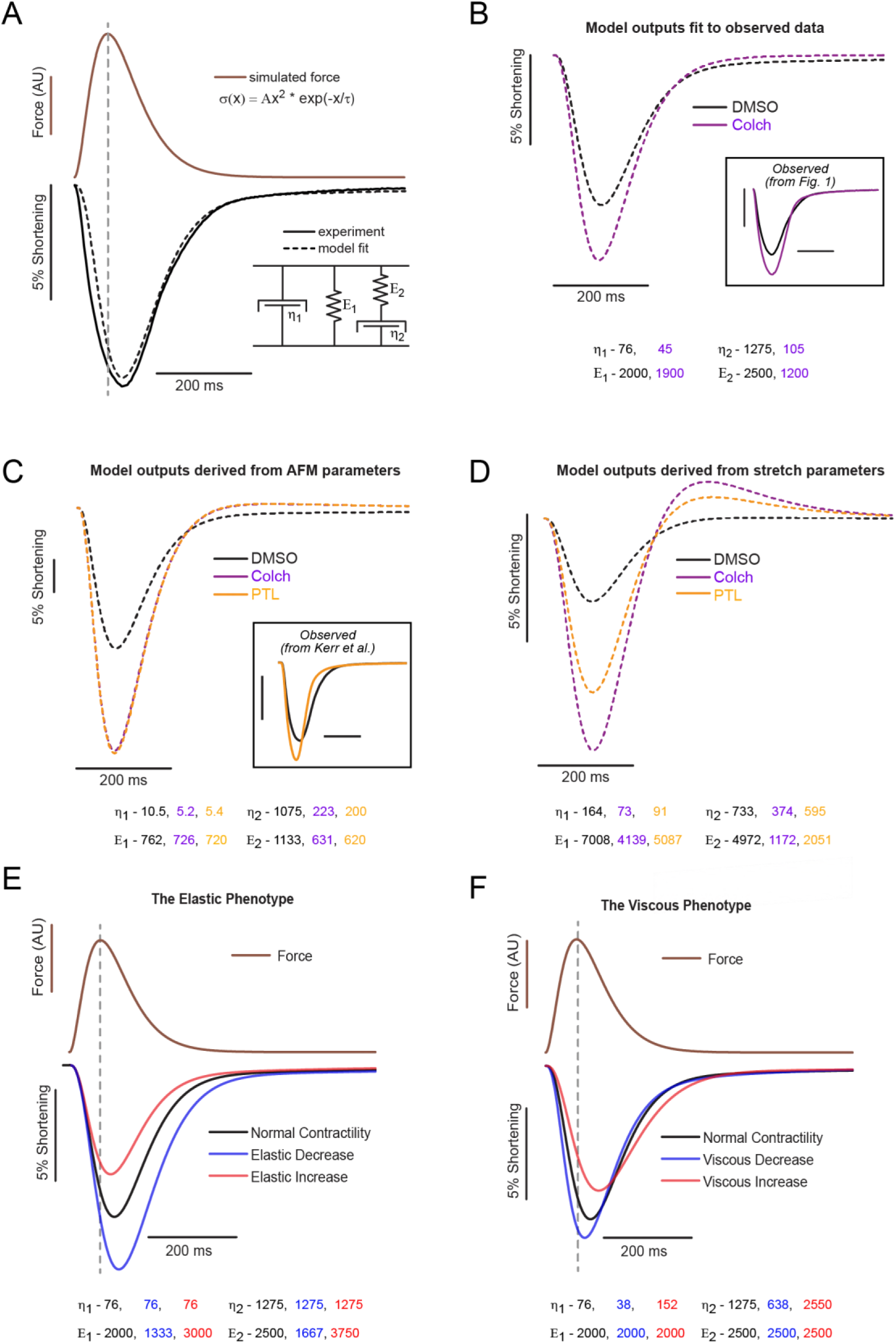
Viscoelastic model of cardiomyocyte contractility. **A**, The stress-strain relationship defined by the VE model (bottom right inset) converts myofilament force (brown) to shortening (black dotted trace). By changing the stiffness of individual viscoelastic elements (values are shown in **B**, bottom), experimentally observed shortening (black solid trace) can be modeled (black dashed trace) by cytoskeletal viscoelasticity. **B**, Model fits (dashed traces) to observed DMSO and colchicine contractility data (boxed inset). Best fit parameters are shown below the plot. **C**, Modeled contractility from viscoelastic parameters derived from transverse indentation studies (Figure 2). Previously observed data for PTL treatment is provided for comparison (inset). **D**, Modeled contractility from parameters derived from tensile tests (Figure 3). **E**, Simulated changes in contractility for myocyte where there is an isolated reduction (blue) or increase (red) in myocyte elasticity. **F**, Simulated changes in contractility where myocyte viscosity has been decreased (blue) or increased (red).

Fitting the VE model to the measured colchicine contractile transient provides a modeled estimate of the viscoelastic changes that occur in response to MT depolymerization (Figure 5B). The best fit values of the VE model to the contractility transients in Figure 1 are shown below Figure 5B. Under identical force, the changes in cell shortening observed with colchicine treatment can be tightly reproduced by reducing the viscoelastic properties of the myocyte (η_1_, η**2**, and E_2_). Simply changing the elastic properties (E_1_) of the myocyte cannot recapitulate the change in both shortening amplitude and velocity (Figure 5E). Thus, the model agrees with experimental observations that increased contractile amplitude and velocity can be imparted by a decrease in myocyte viscoelasticity, but not by elasticity alone.

Using the measured viscoelastic parameters obtained by transverse indentation or axial stretch (Figure 5 C and D, bottom), the VE model qualitatively reproduces the alterations in myocyte shortening observed upon PTL or colchicine treatment. Quantitatively, indentation underestimates where axial stretch overestimates the myocyte stiffness felt during shortening, leading to a respective overestimate and underestimate of the degree of shortening predicted by the model. This disparity suggests that transverse indentation is more sensitive to probing viscoelasticity imparted by the non-sarcomeric cytoskeleton and underestimates the contribution of longitudinal components of the sarcomere such as titin and the myofilaments. This is consistent with fluorescent observation of the MT network deforming extensively during transverse indentation in the absence of sarcomere stretch (Supplemental Movie 1). Conversely, stiffness measured during axial stretch may be augmented by resting myofilament tension that is not anticipated to impact myocyte shortening.

The model also predicts that viscous energy dissipation (damping) is important for regulating the contractile timescale of the myocyte. If the viscoelastic terms are outweighed by elasticity the contractile timescale closely follows the time course of myofilament force generation. Whereas the reduced contractile timescale imparted by viscoelastic reduction indicate that myocyte viscosity is sufficient to impact the contractile timescale of the unloaded myocyte. Consequently, there is likely a balance between elastic restoring forces and viscous damping in regulating myocyte contractility.

The model distinguishes between two contractile phenotypes that can arise from different mechanical underpinnings (Figure 5 E,F). An elastic phenotype, as shown in Figure 5E, is characterized by changes in contractile amplitude proportional to changes in timescale. These changes would be signature of changes in myocyte restoring force (perhaps due to a change in titin/myofilament compliance, whereby an increase in restoring force reduces contractility but accelerates relaxation. Conversely, reducing the stiffness of series elastic elements increases contractility but extends the contractile timescale. On the other hand, viscous changes, as shown in Figure 5F, are characterized by changes in contractile amplitude that are inversely proportional to changes in the contractile timescale. Reducing myocyte viscosity, as is accomplished above by reducing MT-dTyr, enhances myocyte contractility while accelerating the contractile timescale, while increased viscosity reduces contractility but lengthens the contractile timescale. This is consistent with the idea that internal viscosity in the myocyte resists motion in both directions.

## 4. Discussion

Here we find that viscoelastic resistance imparted by microtubules is sufficient to impede cardiomyocyte motion. Myocytes have various mechanisms to regulate elastic and viscous characteristics – for example changes in the stiffness of spring elements can occur by changes in titin isoform expression or post-translational modification (23-25). Our work in conjunction with others suggest that the microtubule network contributes viscosity to the myocyte (6-8). This contribution can be regulated by the density of the MT network or by certain post translational modifications that can alter MT stiffness (acetylation)(26) or cross-linking with other cytoskeletal binding partners (detyrosination)(2, 27). Herein, we show that post-translational detyrosination of the MT network increases myocyte viscosity, and that the pool of detyrosinated MTs largely accounts for the MT contribution to myocyte mechanics.

The experimental measurements and modeling are derived under the assumption of linearity. Muscle tissue exhibits clear non-linear mechanical properties –, thus consideration was taken that both transverse indentation and tensile tests were performed largely in the linear regime, for example straining a cell from a SL of 1.8 to 2.1 μm during diastolic stretch. We feel the main finding of this work - that MT-dependent viscosity impedes the motion of rat cardiomyocytes - is strongly supported by the results, however caution should be taken in extending this finding to different mechanical conditions such as significant preload or afterload, or at different shortening rates and amplitudes. Nevertheless, this study highlights the importance of considering that muscle shortening is not a purely elastic process and is therefore not 100% energetically efficient.

The mechanical role of MTs has been reported to be viscous(8), elastic(9), viscoelastic(7), and not significant(**28**) in cardiomyocytes. Our results help to reconcile these measurements, whereby the measurement direction and strain rate determines how and whether MTs contribute significantly to myocyte stiffness. At slow deformation rates, we find a small or insignificant change in MT-dependent mechanical properties, which is consistent with modest changes observed during previous measurements of resting stiffness (6, 9). At higher strain rates we observe a significant contribution of the MT network that is consistent with a role in myocyte viscoelasticity. Our high-speed deformations are consistent with the strain-rates experienced by the myocyte during the contractile cycle, specifically during early-stage diastolic filling. We complement our mechanical measurements and unloaded shortening measurements with computational modeling that quantitatively supports the idea that alterations in viscosity and viscoelasticity underlie the observed changes in myocyte mechanics.

While previous studies have reported non-significant effects of colchicine on the contractility of healthy rat myocytes, the basal contractile velocity reported was less than half of what we obtain (19). A slower rate of contraction would diminish the viscous resistance of MTs, thus accounting for the lack of effect of MT depolymerization. By the same standard, the slower contractile velocities in larger animal models may render MT resistance largely undetectable unless the MT network is significantly proliferated and detyrosinated, as it is in disease(2, 3, 29), and in agreement with the findings of Cooper et al (10). Thus, while previous reports on MT mechanics at first appear contradictory, many of these findings are quite reconcilable when rate-dependence is taken into consideration.

Our work shows that viscosity plays a significant role modulating the contractility of single unloaded myocytes, but our condition of zero preload will likely give the largest effect of viscosity. The Fenn effect argued against viscoelastic theory by demonstrating that the energy liberated during contraction is equal to the sum of the isometric energy and the work therefore suggesting that muscle is highly elastic(30). However, studies utilizing an isotonic baseline suggest that cardiac muscle exhibits an energetic cost to shorten even under load (30, 31). This may be attributable to viscous dissipation as the decline in efficiency also scales with shortening velocity and would be predicted to impact cardiac function. If viscosity plays a significant role in cardiac contractility, we would anticipate that and increase in viscosity would reduce the work done by the heart during a contractile cycle playing the most significant role during phases of rapid volume change in the heart. This would manifest as reduced area and increased slope during rapid diastolic filling in a pressure-volume loop. Both these characteristics are frequently observed in failing hearts{Warriner:2014bf}, but could be attributable to a number of alternative sources such as decreased force production and/or tissue stiffness.

This study aimed to carefully characterize the MT contribution to mechanics at the single myocyte level. As MTs also play important roles in the formation and maintenance of structures involved in cell-cell and cell-matrix connections, tissue level examinations, while likely more complicated, are a clear next step. Further, as the cell stretch technique used here could be sensitive to both shear resistance and axial tension, tissue studies could preferably isolate axial tension. Previous myocardial strip experiments suggest a viscoelastic contribution of MTs (32), but these results must be replicated and expanded, and detyrosination has not been explored.

In summary, depolymerization or tyrosination of MTs enhances myocyte contractility and contractile kinetics independent of changes in calcium handling. Transverse indentation and axial stretch independently show that MTs contribute to myocyte viscoelasticity by a mechanism that is dependent on post-translational detyrosination. At slow strain rates the steady state stiffness of isolated rat myocytes is not significantly dependent on the MT network, but a progressive contribution to stiffness emerges as the strain-rate of the mechanical test is increased. At deformation rates consistent with cardiomyocyte contractility, myocytes treated with colchicine or PTL are significantly softer, providing a mechanical basis for the increased contractility. Computational dissection of the viscoelastic influence on contractility indicates that the contractile phenotype observed with MT depolymerization/tyrosination can be explained by changes in myocyte viscoelasticity. In sum, the findings indicate that MTs act as cytoskeletal shock-absorbers, which mechanically regulate the amplitude and timescale of myocyte contraction by resisting rapid length changes during both systole and diastole.

## 6. Glossary

AdV: – Adenovirus
BDM: – butanedione monoxime
Colch: – colchicine
DMSO: – dimethylsiloxane vehicle
dTyr: – detyrosinated
E: – elasticity
*ϵ*: – strain
η: – viscosity
MOI: – multiplicity of infection
MT: – microtubule
PTL: – parthenolide
NF: – non-failing
NT: – normal Tyrode’s solution
RCM: – rat cardiomyocyte
SL: – sarcomere length
σ: – stress
TTL: – tubulin tyrosine ligase
VE: – viscoelastic

## 7. Acknowledgements

This work was supported by funding from National Institute of Health (NIH) R01-HL133080 to B.L.P., American Heart Association 17POST33440043 to M.A.C., and by the Center for Engineering MechanoBiology (CEMB) through a grant from the National Science Foundation’s Science and Technology Center program: 15-48571. The procurement of human heart tissue was enabled by grants from NIH (HL089847 and HL105993) to K.B.M. We would like to thank Brandon Kao for assistance with nanoindentation.

---

**Supplementary Table 1:** Values for calcium and contractility.

| Contractility Transient | Control |  | Colchicine |  | P | significance |
| --- | --- | --- | --- | --- | --- | --- |
|  | Mean | SE | Mean | SE |  |  |
| Resting Sarcomere Length ($\mu\text{m}$) | 1.826 | 0.008 | 1.829 | 0.01 | 0.997 | |
| Peak % shortening | 9.38 | 0.507 | 11.94 | 0.651 | 0.003 | ** |
| Departure Velocity ($\mu\text{m/s}$) | -3.50 | 0.264 | -5.06 | 0.374 | 0.001 | *** |
| Return Velocity ($\mu\text{m/s}$) | 2.53 | 0.198 | 3.39 | 0.199 | 0.003 | ** |
| t to peak 90% (ms) | 81.2 | 3.0 | 74.9 | 1.9 | 0.07 |  |
| t to peak 50% (ms) | 34.2 | 1.3 | 30.2 | 1.2 | 0.029 | * |
| t to baseline 90% (ms) | 241.1 | 11.6 | 221.7 | 9.9 | 0.206 |  |
| t to baseline 50% (ms) | 147.4 | 9.2 | 144.8 | 8.2 | 0.836 |  |
| Calcium Transient | Control |  | Colchicine |  | P |  |
|  | Mean | SE | Mean | SE |  |  |
| Amplitude ($F/F_0$) | 6.18 | 0.29 | 6.11 | 0.28 | 0.853 | |
| Time to 50% Rise (ms) | 18.09 | 0.43 | 18.19 | 0.51 | 0.889 |  |
| Time to 90% Rise (ms) | 27.91 | 1.04 | 29.71 | 1.33 | 0.296 |  |
| Time to 50% Decay (ms) | 192.4 | 4.8 | 192.7 | 3.2 | 0.958 |  |
| Time to 90% Decay (ms) | 426.7 | 11.9 | 406.7 | 10.1 | 0.201 |  |

**Supplementary Movie 1:** Nanoindentation of rat cardiomyocyte. AdV-EMTB 3x GFP transfected in culture for 48 h enabled MT imaging during nanoindentation. During indentation MTs deform significantly, while sarcomeres are not stretched. Two color overlay shows initial frame (pre-indentation) in blue and subsequent frames (during indentation) in orange. Colocalization appears white.

